# Interactions of Natural Compounds and Biomolecules with Hepatitis C Virus RNA Untranslated Regions: Exploring Structural Modifications to Advance Pathogenesis Understanding and Antiviral Strategy Design

**DOI:** 10.1101/2025.06.13.659499

**Authors:** Raskar Dhanashri Anil, Sakshi Y. Mastoli, Dipali Barku Dongare, Aman, Lalbiakmawia, Avantika Bhatia, Nidhi Srivastava, Shubhini A. Saraf, Abhishek Dey

## Abstract

Hepatitis C virus (HCV), a 10 kb positive-strand RNA virus, primarily infects liver cells and is a major cause of liver cirrhosis, which can progress to hepatocellular carcinoma. The HCV RNA untranslated regions (UTRs) adopt topologies that serve as binding sites for host proteins and miRNAs, crucial for viral replication and pathogenesis. However, the precise interactors and mechanisms of these interactions remain unclear. This study investigates the architectural landscape of HCV UTRs alone and in the presence of various interactors. Structural analysis reveals stable HCV UTR topologies, but significant alterations occur upon interaction with miRNAs and phytochemicals, potentially critical for viral pathogenicity. Additionally, proteins SRSF7 and AKAP8L are identified as novel interactors, which could have implication in host immune evasion and enhanced virulence. These findings highlight new miRNAs, proteins, and small molecules as interactors of HCV UTRs, offering insights into HCV infectivity and therapeutic targets for antiviral interventions.

---

Hepatitis C is a liver condition caused by the hepatitis C virus (HCV), in which the liver becomes inflamed. The virus can cause both chronic and acute hepatitis, resulting in serious illness, including liver cirrhosis and cancer. HCV belongs to the Flaviviridae family and consists of positive single-stranded RNA ∼10 kb in length. The viral RNA consists of a large open reading frame (ORF) flanked on either side by untranslated regions (UTRs). Both 5’ and 3’ UTRs of HCV RNA are the most conserved regions of viral RNA in the HCV strain.^1, 2^ The ∼340 nucleotide (nt) 5’ UTR consist of heavily structured regions with numerous stem-loop helices.^3^ Domains II-IV in HCV 5’ UTR form Internal Ribosomal Entry Sites (IRES), which act as an entry site for host ribosomes to bind to viral RNA and initiate translation in cap-independent manner.^4^ It is also implicated in interacting with miR-122, the binding of which is critical for viral replication.^5^ The 3’ HCV UTR is ∼ 250 nt with a long poly U stretch of about 90 nt long.^6^ It has been found to enhance viral translation, possibly stabilizing the 5’ UTR region through long-range interactions.^7^ Additionally, these UTRs are also known to interact with many host proteins thus enhancing viral infection and pathogenicity. For instance, the HCV 3’X region (3’ UTR terminal region) was found to interact with the host ribosomal proteins thus enhancing the viral RNA translation possibly by directing the ribosomes to the IRES of 5’ UTR.^8^ Furthermore, nucleotide modifications in these regions allow them to mimic host RNA, effectively evading detection by the host immune system. Due to their crucial role in viral infection, HCV UTRs are the focus of extensive research and are continually targeted for novel antiviral drug development. In this study, we investigate the conformational flexibility of HCV UTRs regions in the presence of naturally occurring small molecules. Our study indicates flexible HCV UTR topology that can be destabilized in the presence of small molecules. HCV UTR also acts as binding sites of host microRNA (miRNA)^9^. The current work identifies novel host miRNAs capable of interacting with the HCV UTR, inducing structural modifications. Current study also identifies novel host proteins as potential interactors of HCV UTR which can help in their nucleotide modifications and enhance viral pathogenesis.

To analyse the secondary structure, HCV 5’ UTR (178nt-341nt) and 3’ UTR (9545nt-9646nt) were *in vitro* transcribed (see materials and methods). Circular Dichroism (CD) analysis of the constructs revealed a sharper positive peak for the 5’ UTR within the range of 260–280 nm, with a midpoint near ∼265 nm, and a weak CD signal after 290 nm. Additionally, a distinct negative peak was observed around 200–220 nm, with a midpoint at ∼210 nm (**Figure 1A**). These spectral features suggest that the HCV 5’ UTR adopts an A-form topology, characterized by extensive base pairing that stabilizes the RNA conformation.^10-12^ The CD spectra of the 5’ UTR also indicate the potential presence of a pseudoknot structure, as suggested by similarities with previously reported spectra for unrelated RNAs.^13^ To further investigate the secondary and tertiary structures of the HCV 5’ UTR, various RNA folding algorithms were employed. Vienna RNAfold predicted a highly structured RNA consisting of five stem-loop helices (Stem I, Stem II, Stem III, Stem IV, and Stem V) with internal loops, bulges, and apical loops of varying sizes **(Figure 1B**). In contrast, IPknot predicted an alternate topology featuring five distinct stem-loop helices with potential long-range nucleotide interactions, resulting in pseudoknot formation **(Figure 1C)**.

**Figure 1:**
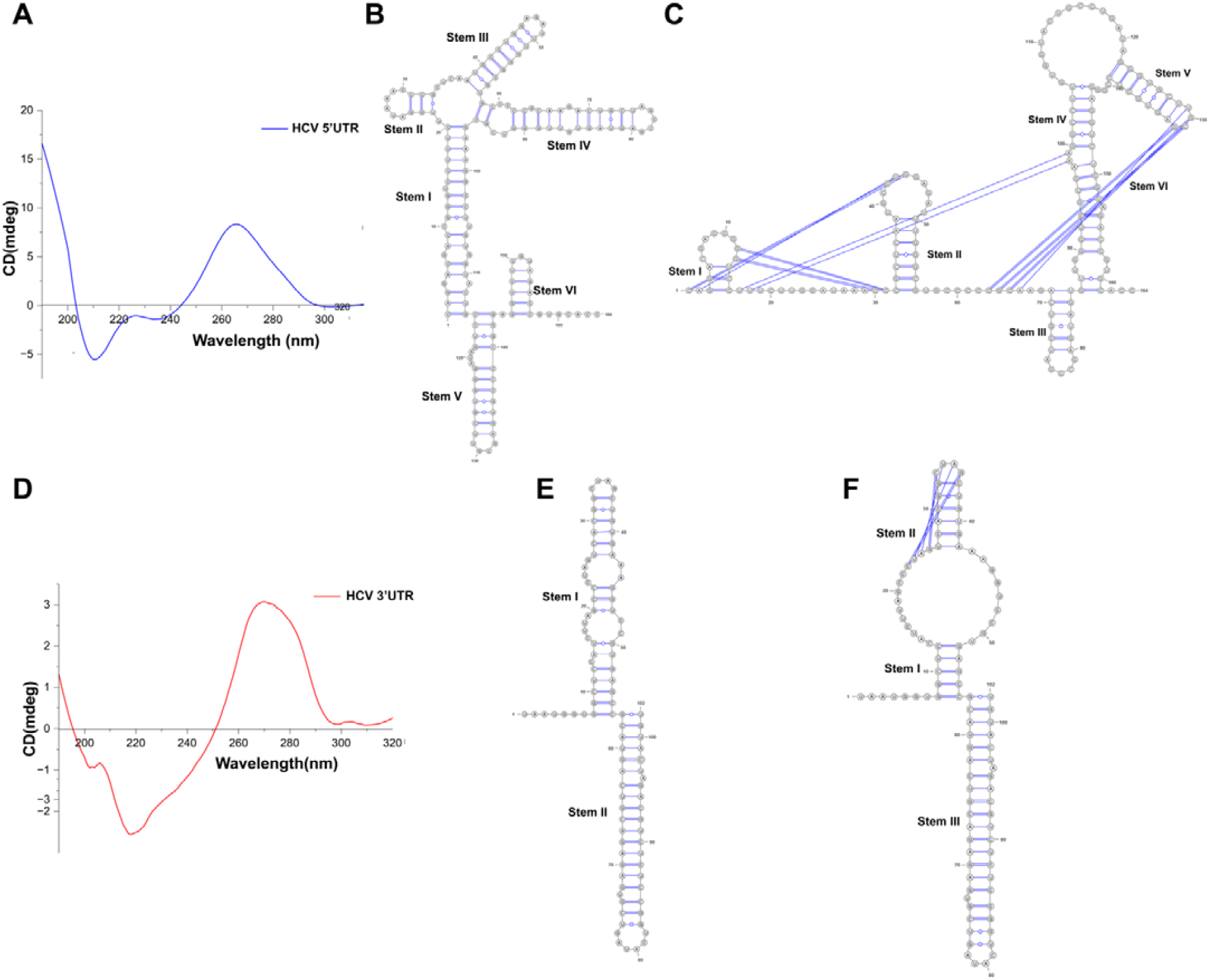
Architecture of HCV 5’ and 3’ UTR reveals stable stem-loop helices with tertiary interactions. Secondary structure of HCV 5’ UTR determined through (A) Circular dichroism ViennaRNA (C) IPknot. Secondary structure of HCV 3’ UTR determined through (D) Circular dichroism (E) ViennaRNA (F) IPknot.

The CD spectra of the HCV 3’ UTR exhibit distinct features compared to those of the 5’ UTR. A positive peak in the 260–280 nm range and a negative peak in the 200–220 nm range indicate an A-form RNA structure **(Figure 1D)**. However, the broader and less intense peaks suggest reduced base pairing, consistent with a more open RNA conformation **(Figure 1D)**. *In silico* folding analysis using ViennaRNA predicts a two-stem helical structure (Stem I and Stem II) for the HCV 3’ UTR, both terminating in apical loops **(Figure 1E)**. Additionally, Stem I includes two internal bulges **(Figure 1E)**. Predictions from IPknot also reveal a two-stem helical topology but identify a large internal loop in Stem I. Nucleotides 23–26 engage in tertiary interactions with the apical loop, resulting in a pseudoknot conformation **(Figure 1F)**. Together, the CD analysis and *in silico* predictions indicate the presence of folded and stabilized conformations in HCV UTRs.

The 5′ UTR of HCV contains an Internal Ribosomal Entry Site (IRES) that facilitates cap-independent translation initiation of the viral RNA by enabling host ribosome recruitment. Previous cryo-EM^14^ and in solution small-angle X-ray scattering (SAXS)^14^ studies have demonstrated the folded conformational topology of the HCV IRES. Complementing these findings, our circular dichroism (CD) and computational analyses of the HCV UTRs confirm the presence of stable stem-loop helices.

HCV 3’ UTR construct used in our study consists of the region downstream of the poly(U). Incidentally, this region is also consists of 3’X region which is also essential for IRES dependent translation.^15^ Previous RNA-SHAPE study have identified a stable 3 stem-loop helical conformation post poly(U) region for HCV 3’ UTR, which is crucial for interacting with 40S ribosome to initiate IRES dependent translation.^7^ Our CD and *in silico* analysis have identified HCV 3’ UTR as a least stable conformation with two stem helices. Moreover, our study have also uncovered additional long-range base pairing, indicative of pseudoknot formation in both 5’ and 3’ UTR. However, their role in viral proliferation and pathogenicity remains to be elucidated.

To evaluate the impact of small molecules on the secondary structure of the HCV UTR, we performed circular dichroism (CD) analysis of the HCV 5’ UTR and 3’ UTR in the presence of naturally occurring compounds, including phytic acid, caffeine, and nicotine. Phytic acid, known for its potent antioxidant and anti-cancer properties^16, 17^, has an unexplored role in HCV-derived hepatocellular carcinoma. Our CD analysis revealed that phytic acid induces significant changes in the secondary structure of both the HCV 5’ and 3’ UTRs (**Figures 2A, 2D**). Notably, phytic acid exerts a more pronounced effect on the 5’ UTR, leading to a complete loss of secondary structure **(Figure 2A)**. For 3’ UTR, phytic acid reduces peak intensity and sharpens the peaks, suggesting the disruption of existing base-pairing interactions and the formation of new ones **(Figure 2D)**. Caffeine has demonstrated beneficial effects in reducing liver inflammation and lowering the risk of hepatocellular carcinoma in patients with chronic HCV infections.^18, 19^ CD spectral analysis of the HCV 5’ UTR with caffeine revealed loss of secondary structure **(Figure 2B)**. In contrast, caffeine’s effect on the HCV 3’ UTR reduces its peak intensity **(Figure 2E)** indicating diminished base pairing interactions. Like caffeine, Nicotine has also shown antiviral effect by inhibiting the proliferation and replication of severe acute respiratory syndrome coronavirus 2 (SARS-CoV-2).^20^ 30 µg of nicotine resulted in the loss of CD spectra for HCV 5’ UTR and decrease in peak intensity of HCV 3’ UTR **(Figure 2C and 2F)**, indicating changes in HCV UTR architecture **(Figure 2C and 2F)**. To assess the binding affinity of these compounds we also performed molecular docking studies. Our docking analysis identifies these small compounds as potent interactors of HCV UTRs with high binding affinity *(data not shown)*.

**Figure 2:**
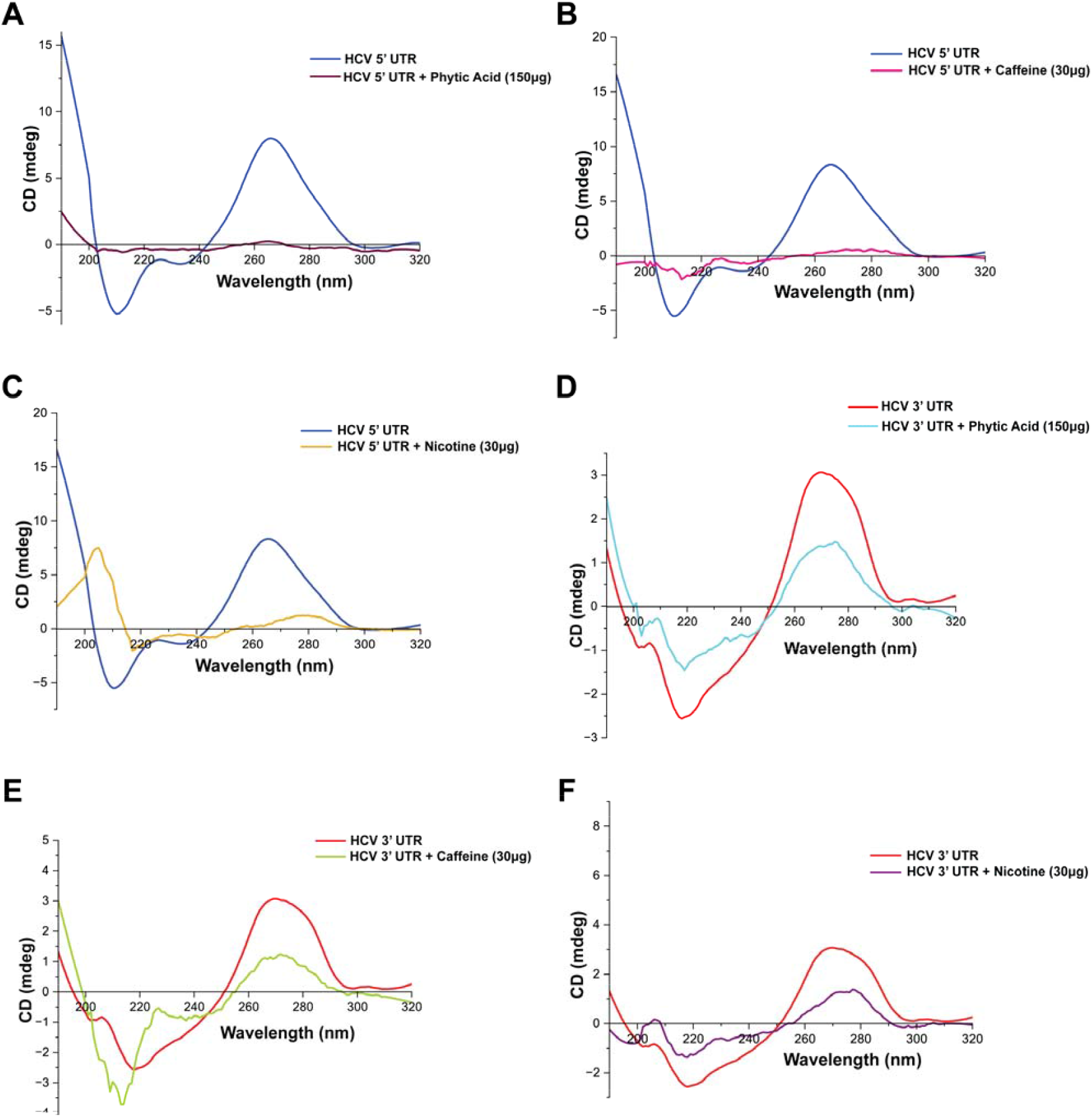
Disruption of HCV UTRs architecture in the presence of natural compounds. CD spectra of HCV 5’ and 3’ UTRs respectively in the presence of (A) and (D) 150 µg of Phytic acid, (B) and (E) 30 µg of Caffeine, and (C) and (F) 30 µg of Nicotine.

Among the three compounds tested, caffeine is the only one known to exhibit protective effects on the liver against hepatitis C virus (HCV) and hepatocellular carcinoma.^19^ Our study is the first to demonstrate the direct impact of caffeine on the architecture of the HCV UTR, showing significant structural modulations. While there is no direct evidence of phytic acid’s antiviral properties, its antimicrobial activity and anti-biofilm effects are well established.^21^ Circular dichroism (CD) analysis from our study also revealed notable alterations in the HCV UTR structure in the presence of phytic acid. The functional implications of these structural changes induced by these natural compounds remain to be explored. In contrast, nicotine is known to have detrimental effects on the liver, particularly in individuals with a history of smoking and HCV infection.^22^ However, interestingly, our study reveals disruption of the HCV UTRs architecture in the presence of nicotine. The natural compounds disrupting the IRES architecture of the HCV 5’UTR, warranting further investigation into their impact on HCV proliferation and infectivity. This study highlights the potential of natural small molecules as anti-HCV agents.

Chronic HCV infections can lead to the upregulation of certain miRNA and long non coding RNAs (lncRNAs) that can have oncogenic effects resulting in the development of HCC or liver cancer.^23^ H19 lncRNA and its derived miR-675 has been implicated in development and progression of many cancers,^24^ including HCC.^2^ Overexpressed miR-675 downregulates the expression of Twist1 and retinoblastoma (Rb) protein thus enhancing the proliferative capacity of liver cells and intensifying the progression of HCC.^2^ Similarly, miR-331-3p is known to promote the development of HCC by enhancing cellular proliferation and metastasis. Overexpressed miR-331-3p stimulates protein kinase B (AKT), thus enhancing epithelial mesenchymal transition (EMT) in HCC.^25^ To assess whether these highly expressed miRNA can interact with HCV UTRs, we further performed CD analysis on these miRNAs alone and in conjugation with HCV UTRs. CD spectra obtained for miR-675 in current study have a sharp and intense positive peak in the 260–280 nm range and a negative peak in the 200–220 nm range indicative of A-form RNA stabilized by base-pairing interactions. Our earlier study on miR-675 determined it as a single stem-loop helical architecture (**Figure 3A, inset**)^24^ thus supporting current observations (**Figure 3A**). Upon conjugation of miR-675 with HCV 5’ UTR, very subtle changes in the CD spectra of conjugated RNA was observed, indicative of weaker interactions between these two molecules (**Figure 3B**). However, the CD analysis of HCV 3’ UTR conjugated miR-675 with reveals a distinct pattern where the peak intensity of coupled RNA is sandwiched between the individual peaks of miR-675 and HCV 3’ UTR (**Figure 3C)**. Moreover, this spectra was also sharper than what obtained for HCV 3’ UTR alone (**Figure 3C**). This indicates the opening of the secondary structure of both RNA and formation of new base pairs between the two to form a hybrid RNA.

**Figure 3:**
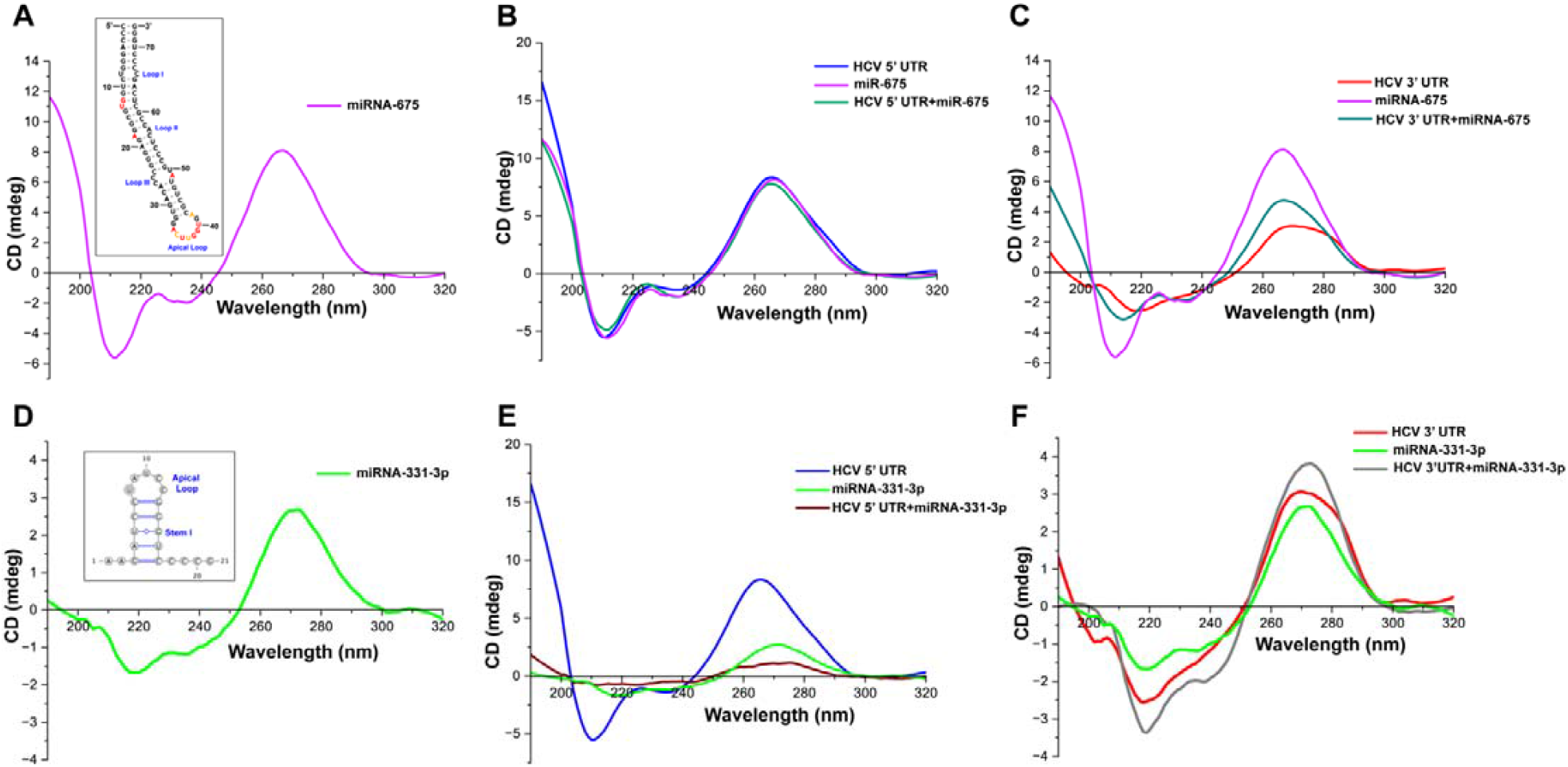
Interactions between HCV UTRs and miRNA through structural modulations. CD spectra of (A) miR-675, as stem-loop helix in inset^24^ (B) HCV 5’ UTR and miR-675, (C) HCV 3’ UTR and miR-675. CD spectra of (D) miR-331p-3p, as stem-loop helix in inset, (E ) HCV 5’ UTR and miR-331-3p, and (F) HCV 3’ UTR and miR-331-3p.

CD spectra obtained for miR-331-3p was similar to the spectra obtained for miR-675 (**Figure 3D**), however, the negative spectral intensity at the range of 200-240 nm was found to be less intense (**Figure 3D**). This indicates the presence of open conformation of miR-331-3p. Indeed the viennaRNA fold predicted a lone stem-loop helical conformation of miR-331-3p with the stem I being stabilized by five base pairing interactions (**Figure 3D**, inset). Spectral analysis of miR-331-3p in conjugation with both HCV 5’ UTR and HCV 3’ UTR reveals distinguishable secondary structure features for hybrid RNA. Peak intensity of conjugated miR-331-3p and HCV 5’ UTR was found to be less intense when compared to the spectra obtained for alone HCV 5’ UTR (**Figure 3E**). It seems that both miR-331-3p and HCV 5’ UTR lose their secondary structure when they are co-existing. However, CD spectra obtained for coupled miR-331-3p with HCV 3’ UTR reveals a peak with more intensity at 260-280 nm suggesting strong interactions between both HCV 3’ UTR and miR-331-3p. Though there is no known earlier evidence for the direct interactions between these two miRNAs with HCV UTRs, our previous co-folding analysis between miR-675 and its parental H19 lncRNA does revealed the intricate feedback regulatory mechanism exerted by both RNA on each other.^24^ This study identifies direct interactions between miRNAs and HCV UTRs that modify their architecture, potentially enhancing viral proliferation and pathogenicity critical to HCC progression.

RNA viruses are known to exploit the host’s protein machinery to support their replication and propagation. In the case of HCV, chemical modifications of its RNA nucleotides play a crucial role in regulating its life cycle. The N^6^-methyladenosine (m6A) modification of HCV RNA suppresses the host’s immune response by disrupting RIG-I sensing activity, thereby downregulating interferon signalling within the host cell.^26^ This makes the virus stealthier to effectively evade the host immune response. RNA modifications are mediated by various host proteins, including writers, erasers, and readers of modified nucleotides. However, the specific proteins capable of interacting with and modifying the HCV UTR remain unidentified. Using RBPsuite, potential proteins were screened for their ability to either modify the HCV UTR or recognize and bind to these modifications. Among the candidates, Serine/Arginine-rich splicing factor 7 (SRSF7) and A kinase anchoring protein 8-like (AKAP8L) emerged as potential interactors of the HCV UTRs. SRSF7 is an RNA-binding protein involved in the regulation of mRNA splicing and export and has been implicated in the progression of cancer.^27^ SRSF7 interacts with YTHDC1, an N6-methyladenosine (m6A) reader protein implicated in hepatocellular carcinoma progression.^28^ To validate the interaction between SRSF7 and the Hepatitis C Virus (HCV) untranslated region (UTR), a super shift assay was performed using HCV 5′ UTR RNA, HEK 293 cell lysate, and an anti-SRSF7 antibody. A higher-molecular-weight band corresponding to the HCV 5′ UTR RNA complexed with the anti-SRSF7 antibody was observed in the NATIVE gel, indicating interactions between SRSF7 and the HCV 5′ UTR (**Figure 4A**). The interaction between the HCV 5’ UTR and SRSF7 was investigated through intermolecular interaction analysis using HADDOCK 2.4.^29^ Our analysis identified ionic interactions between the basic residues of SRSF7 and the acidic regions of the HCV 5’ UTR as the primary driving force behind the interaction between the two molecules (**Figure 4B**). Similarly, interaction between the HCV 3′ UTR and AKAP8L was also identified using super shift assay with an anti-AKAP8L antibody (**Figure 4C)**. Additional HADDOCK analysis revealed a similar interaction pattern between AKAP8L and the HCV 3’ UTR, akin to that predicted for SRSF7 and the HCV 5’ UTR (**Figure 4D**). AKAP8L is known to enhance the replication of HIV^30^ by interacting with other host proteins and viral RNA. It is also known to regulate the activity of histone deacetylase thus controlling the epigenetic modification of the genome.^31^ While the role of AKAP8L protein in HCV proliferation and disease progression remains unclear, its interaction with HCV 3’ UTR regions indicates the possible chemical modifications of HCV UTR critical for viral proliferation and pathogenesis. This study however identified SRSF7 and AKAP8L as novel interactors of HCV UTRs and opened up new possibilities of exploring both as a potential target for anti-HCV therapy.

**Figure 4.**
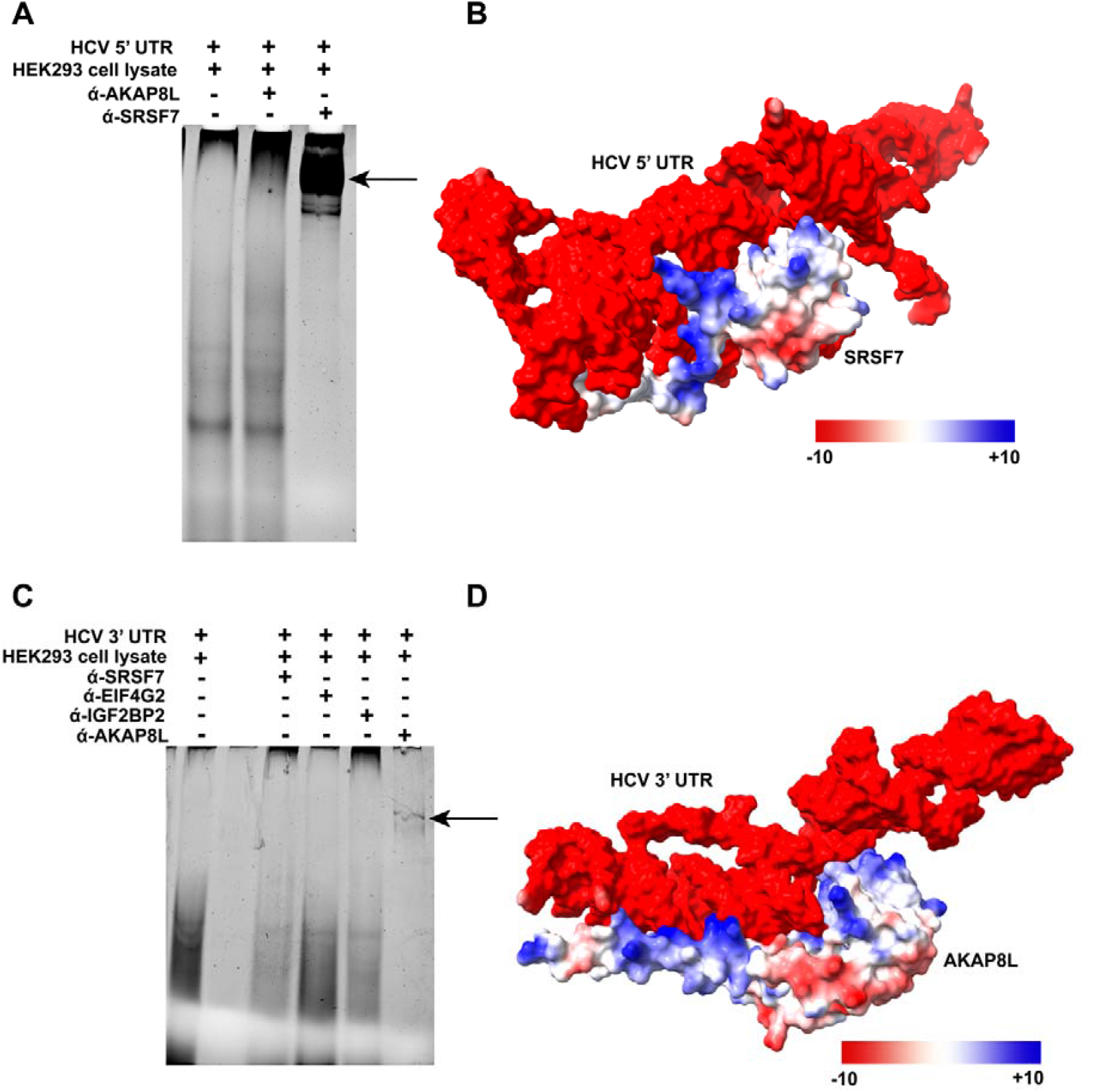
SRSF7 and AKAP8L proteins bind to HCV UTRs. Interaction of SRSF7 with HCV 5’ UTR assessed through (A) gel super-shift assay, arrow indicating shift in HCV 5’ UTR band in gel and (B) ionic interactions predicted using HADDOCK. Interaction of AKAP8L with HCV 3’ UTR assessed through (C) gel super-shift assay, arrow indicating shift in HCV 3’ UTR band in gel and (D) ionic interactions predicted using HADDOCK.

## Materials and Methods

### *In Vitro* Transcription of HCV UTRs

For conformational analysis, HCV 5’ UTR (178nt-341nt) and 3’ UTR (9545nt-9646nt) with T7 promoter regions at 5’ end were synthesized from sigma oligos as double stranded DNA. These regions were PCR amplified and were further used as a template for *in vitro* transcription reaction using T7 high yield RNA kit (New England Biolabs). The synthesized RNA was DNase-treated (TURBODNase), purified using RNA cleanup kit (Geneaid) and quantified with nanodrop.

## Circular Dichroism Spectroscopy

For CD analysis 32μg of HCV 5’ UTR and 36μg of HCV 3’ UTR was diluted in phosphate buffer (pH 8.3) to the final volume of 500µl. For CD analysis with naturally occurring small molecules, 32μg HCV 5’ UTR and 36μg of HCV 3’ UTR were incubated with 150μg of phytic acid, 30 μg of caffeine and nicotine respectively for 1 hr at 37□. For interaction analysis between HCV UTRs and miRNAs, 40μg HCV 5’ UTR and 45μg HCV 3’ UTR were mixed with 41μg of miR-675 and 28μg of miR-331-3p respectively. The entire reactions were incubated at 37□ for 1 hour. All CD measurements were taken in a JASCO CD spectrophotometer with a scan speed of 50 nm/min, and an excitation bandwidth of 1.00 nm. Integration period of 1s were used to capture spectra over a wavelength range of 190 to 320 nm. OriginPro 2024 was used to construct the CD spectra. A total of 3 independent (n=3) samples for each reaction were used for spectra measurement.

### *In silico* secondary structure predictions

For *in silico* secondary structural predictions of the RNA constructs, we utilized ViennaRNA^32^ and Ipknot,^33^ both of which are widely recognized for their robust algorithms in RNA structure prediction. These tools were applied to both RNA constructs under study, and the predictions were performed using their default parameter settings.

## RBPsuite to identify protein interactors of HCV UTRs

To identify potential interacting partners and protein-binding sites on HCV UTRs, searches were conducted using RBPsuite^34^ with default settings. HCV UTR was input as a linear RNA with the general model selected as the prediction framework. Protein-binding sites are represented by binding scores ranging from 0 to 1, where higher scores indicate a greater likelihood of an RBP site within the RNA sequence.

### Gel super shift assay

For gel supershift assay 32μg of HCV 5’ UTR and 36μg of HCV 3’ UTR of HCV UTR was incubated with HEK293 cell lysate (∼1.5 million cells) for 1 hr at 37□. After incubation, 2ul of anti-SRSF7 (E-AB-66923, Elabscience) and anti-AKAP8L (DF12818, Affbiotech) antibodies were added to the reaction mixture and further incubated for 1 hr at 37□. All the samples were finally loaded into 8% NATIVE gel and separated using 1X Tris-Borate EDTA buffer. Gels were post-stained with EtBr and images were captured using UV shadowing using gel documentation system (BioRad).

### HADDOCK run

HADDOCK is a flexible, information-driven approach for modeling biomolecular complexes. Using HADDOCK 2.4, interactions between the Hepatitis C virus (HCV) 5’UTR and SRSF7, and the 3’UTR and AKAP8L, were modeled. Active residues for SRSF7 (PDB ID: 2HVZ) (1-101), 5’UTR (15-164), AKAP8L (401-543), and 3’UTR (1-102) were selected, while passive residues were auto-assigned by the software. The 5’UTR-SRSF7 docking yielded six clusters, with Cluster 6 having the best score (Z-score: -2.1). Similarly, the 3’UTR-AKAP8L docking also produced six clusters, with Cluster 6 scoring highest (Z-score: -1.4).

### Molecular Docking

The RNA structure and ligand were prepared in PDB format and edited using AutoDockVina 1.5.7 by removing water molecules, ions, and other unnecessary components. Missing hydrogen atoms were added, and MGLTools was used to convert both structures to PDBQT format. Grid box parameters were set, with the docking box coordinates saved for the HCV 5’ UTR (center: 3.078, -7.226, -2.762) and 3’ UTR (center: -53.480, -33.823, 35.463). After docking binding poses were visualized using Discovery Studio Visualizer.

## ABBREVIATIONS

HCV: Hepatitis C Virus
UTR: Untranslated Regions
HCC: Hepatocellular carcinoma
lncRNA: Long noncoding RNA
miRNA: microRNA
CD: Circular Dichroism.

## AUTHOR INFORMATION

### Author Contributions

^‡^R.D.A, S.Y.M, and D.D.B contributed equally to this work. R.D.A, S.Y.M, and D.D.B performed the experiment, analyzed the data, and reviewed the manuscript; R.D.A performed molecular docking studies; A. helped with some CD experiments; L. performed HADDOCK studies, A.B. helped with gel supershift studies, N.S. reviewed the manuscript, S.A.S, reviewed and edited the manuscript, A.D. conceptualized the study, analyzed the data, wrote the original draft, reviewed and edited the manuscript. All authors have given approval to the final version of the manuscript.

### Notes

The authors declare no competing financial interest.

## Acknowledgments

The authors would like to acknowledge the Department of Biotechnology, Department of Pharmaceutics, Central Instrumentation Facility (CIF), and Central In-Vitro Facility, NIPER Raebareli, for constant support received while conducting the study and preparation of this manuscript. Dr. Abhishek Dey acknowledges the Department of Biotechnology, Govt. of India, for the Ramalingaswami Re-entry Fellowship (BT/RLF/Re-entry/02/2021). R.D.A, S.Y.M, D.D.B, A., L., and A.B. acknowledges the Department of Pharmaceuticals, Ministry of Chemicals and Fertilizers, Govt. of India for providing fellowship assistance. This article bears NIPER-R communication number 708.

